# Cyclic di-GMP interact with putrescine via a PilZ domain receptor YcgR

**DOI:** 10.1101/2023.03.05.531214

**Authors:** Weihan Gu, Yufan Chen, Zhongqiao Chen, Huagui Gao, Congcong Xie, Lian-hui Zhang, Lisheng Liao

**Affiliations:** Guangdong Laboratory for Lingnan Modern Agriculture, Guangzhou 510642, China; Guangdong Province Key Laboratory of Microbial Signals and Disease Control, Integrative Microbiology Research Center, South China Agricultural University, Guangzhou 510642, China; Research Center of Chinese Herbal Resource Science and Engineering, Key Laboratory of Chinese Medicinal Resource From Lingnan (Guangzhou University of Chinese Medicine), Ministry of Education, Joint Laboratory of National Engineering Research Center for the Pharmaceutics of Traditional Chinese Medicines, Guangzhou University of Chinese Medicine, Guangzhou, 510006, People’s Republic of China

**Keywords:** *Dickeya oryzea*, c-di-GMP, putrescine, bacterial motility, YcgR

## Abstract

The cell motility is one of the key pathogenic factors that contribute to the virulence of *Dickeya oryzea,* which is a prevalent bacterial pathogen capable of infecting a range of crops and plants. We showed recently that the bacterial second messenger c-di-GMP, and the putrescine-mediated quorum sensing (QS) system, are both involved in the regulation of the bacterial motility in *D. oryzea* EC1. In this study, we set to determine whether and how there two signaling mechanisms work together to modulate the bacterial motility. The results showed that the second messenger signaling system interacts with the putrescine QS system via the c-di-GMP receptor YcgR, which could promote the activity of SpeA, the rate-limiting enzyme in the putrescine biosynthesis pathway, thereby increasing the intracellular putrescine levels. However, it was shown that this facilitative effect could be inhibited by c-di-GMP molecules. In addition, we demonstrated the dominance of c-di-GMP over putrescine in the regulation of bacterial motility. The findings from this study provide the first insight into the interaction between c-di-GMP and putrescine in bacteria and provide a valuable reference for the study of intracellular second messenger system and polyamine-mediated quorum sensing system in other bacteria.

**Importance:** *Dickea oryzea* is a major bacterial pathogen capable of infesting many plants and crops, causing significant economic damage to rice and banana production especially. Bacterial motility is a key pathogenic factor of *D. oryzea* to compete for food resources and infect their host species, which is negatively regulated by c-di-GMP and positively regulated by putrescine, respectively. However, the connection between c-di-GMP and putrscine in regulating the motility of *D.oryzea* is not understood. Here we revealed the link and the mechanism of interaction between them, showing that c-di-GMP interact with putrescine via a receptor of c-di-GMP. The significance of our research is in providing the first insight into the interaction between c-di-GMP and putrescine and the methods and experimental designs in our study will provide a valuable reference for subsequent studies on the link between c-di-GMP and putrescine in other bacteria and even the regulatory mechanisms of complex bacterial motility networks, respectively.

## Introduction

Cell motility is an important mechanism of numerous bacterial pathogens for competing food resources and infecting their host organisms. To survive in harshly environmental conditions, bacteria evolved mechanisms allowing them to switch rapidly between planktonic motile and sessile non-motile forms in response to environmental changes. The state of attachment facilitates the adherence of pathogenic bacterial cells to the host surface and enhances the immune response of the host organisms (1, 2), whereas the planktonic state, represented by strong motility, is not only important for the successful invasion of the host, avoidance of host defense responses, colonization and release of pathogenic factors in the host, but is also essential for the successful formation of biofilm on the surface (3–5). Bacterial motility is normally closely linked to chemotaxis, and the combination of the two enables bacteria to detect and pursue nutrients and to reach and maintain their preferred colonial ecological niche (6). Bacterial motility is mainly mediated by the flagella and controlled by the number of flagella, the direction of rotation and also the speed of rotation (7). The research progress over the years have demonstrated that the bacterial cell motility is an important pathogenic factor, and bacteria with impaired motility are significantly less pathogenic than their wild type isolates (8, 9).

Polyamines are a group of biologically active polycationic fatty compounds found in a wide variety of living organisms, the most common polyamines in microorganisms are putrescine, spermidine and spermidine (10). Putrescine is the central product of the polyamine synthesis pathway. In bacteria, there are two main pathways for the synthesis of putrescine, the ornithine pathway and the arginine pathway. In the ornithine pathway, ornithine decarboxylase (SpeC) catalyzes the formation of putrescine from ornithine. In the arginine pathway, arginine decarboxylase (ADC), encoded by the *speA* gene, catalyzes the conversion of arginine to agmatine, which then converts into putrescine by agmatinase (11). Spermidine and spermine are produced by the reaction of putrescine and aminopropyl, catalyzed by spermidine synthase and spermine synthase, respectively (12, 13). In general, polyamines are associated with many cellular processes in bacteria, such as growth under anaerobic conditions, iron carrier synthesis, peptidoglycan synthesis, biofilm formation, bacterial motility, resistance to oxidative stress and the antibiotics (14–18). In recent years, there has been increasing evidence that polyamines are an intraspecies and interkingdom cell-cell quorum sensing signal that not only play a regulatory function in intercellular communication between bacteria, but are also involved in transboundary communication between hosts and pathogens (8, 19, 20). Among them, putrescine is primarily involved in regulation of the bacterial motility and biofilm formation (27–29). However, the molecular mechanism of putrescine in modulation of bacterial motility and biofilm formation remains poorly understood.

Cyclic di-GMP (c-di-GMP) is a widespread second messenger in bacterial species, whose homeostatic is controlled by diguanylate cyclases (DGCs) responsible for c-di-GMP biosynthesis, and phosphodiesterase (PDEs) involved in c-di-GMP degradation. Specifically, DGCs have a GGDEF domain and PDEs contain either an EAL or a HD-GYP domain (21–23). The second messenger has been shown to regulate a number of physiological processes, such as biofilm formation, motility, virulence, cell differentiation and multiplication through various downstream receptors (24, 25). Among them, the PilZ structural domain proteins constitute the most widely distributed c-di-GMP receptor family, which and can be classified into the single PilZ structural domain class, YcgR class, BcsA class, and Alg44 class (26). YcgR was one of the first PilZ structural domain proteins to be discovered and studied, which consists of the N-terminal YcgR domain and the C-terminal PilZ domain. YcgR binds c-di-GMP molecules via the PilZ domain and then regulates bacterial biological functions through protein-protein interactions or protein-DNA interactions (26). Previous study have shown that in *E. coli*, c-di-GMP promotes the interaction of YcgR with the flagellar proteins FilG and FilM to inhibit bacterial motility by modulating the direction and rate of flagellar rotation (27). In *Klebsiella pneumoniae*, c-di-GMP promotes the binding of MrkH, a YcgR-class protein, to the promoters of *mrkHI* and *mrkA*, which encoding the transcriptional regulators that regulate type 3 fimbriae and biofilm foramtion and the major structural component protein of type 3 fimbriae(28, 29), respectively, through its PilZ structural domain (30).

*Dickeya oryzea* is the causative agent of bacterial foot rot disease in rice crops, leading to severe economic losses (31, 32). The pathogen was previously known as *Erwinia chrysanthemi* pv. *zeae* but was reclassified into the new genus *Dickeya* in 2005 (33). *D. oryzea* is capable of producing a range of pathogenic factors, including the phytotoxin zeamines (34), extracellular enzymes (35), and type III effectors (36, 37). Bacterial motility is a secondary virulence determinant of *D. oryzea* that plays a vital role in its colonization and invasion on rice seeds (38–40). Recently, several regulatory mechanisms which modulate bacterial motility have been unveiled in *D. oryzea* EC1 (41), including an acyl-homoserine lactone (AHL) QS system (32), a polyamine-mediated QS and pathogen-host communication system (8), and the c-di-GMP signaling system (9). Among them, putrescine positively regulates the motility of strain EC1 (8), and the accumulated c-di-GMP negatively regulates the bacterial motility through its downstream receptor protein YcgR (9).

To explore the possible connection between putrescine and c-di-GMP signaling systems in the regulation of *D. oryzea* motility, we tested the cellular level of putrescine in *D. oryzea* wild-type strain EC1 and its mutants defective in c-di-GMP degradation and a functional YcgR receptor, respectively. The results showed that the putrescine level was significantly reduced in these mutants compared to the wild-type strain EC1. *In vitro* protein interaction and *in vivo* bacterial two-hybrid assays revealed a direct interaction between YcgR and SpeA, a key rate-limiting enzyme of the putrescine synthesis pathway. Subsequently, we demonstrated that YcgR could promote the arginine decarboxylase enzymatic activity of SpeA to increase agmatine production, however, this activity could be inhibited by adding c-di-GMP molecules. Furthermore, we demonstrated that c-di-GMP occupied a more dominant role in the regulation of bacterial motility phenotype than putrescine. To our knowledge, this is the first report on the interaction between the putrescine-mediated host-pathogen quorum sensing system and the c-di-GMP signaling pathway.

## Results

### Putrescine and c-di-GMP play opposite roles in regulation of *D. oryzea* swimming motility

To explore the potential cross-talking between Putrescine and c-di-GMP signaling pathways, which were shown previously associated with the regulation of the bacterial motility (10, 11), we conducted experiments to compare the regulatory patterns of these two signaling mechanisms in modulation of the bacterial motility under the same experimental conditions. Null mutation of *speA* in *D. oryzea* strain EC1, which encodes a rate-limiting enzyme in the putrescine synthesis pathway, decreased the swimming motility by about 30% compared to the wild type EC1 (Fig. 1A). Exogenous addition of 0.1 mM putrescine could rescue the swimming motility of Δ*speA* (Fig. 1A), validating the role of putrescine in regulation of the bacterial swimming motility. This was further confirmed by deletion of the two putrescine-specific transporter genes, i.e., *potF* and *plaP* (10), in the mutant Δ*speA*. The resultant mutant Δ*speA*Δ*potF*Δ*plaP* showed much reduced swimming motility, which was about 40% of wild type level and could not be restored by exogenous addition of putrescine (Fig. 1A). The second messenger c-di-GMP seemed to exert much stronger influence on the *D. oryzea* motility than putrescine. When all the c-di-GMP synthase coding genes were knocked out in strain EC1, the mutant 15ΔDGC showed drastically enhanced swimming motility, about 480% of the wild type level. Conversely, the mutant 7ΔPDE with all the genes encoding c-di-GMP-degrading enzymes being deleted became nonmotile (Fig. 1B). Complementation of the mutants 15ΔDGC and 7ΔPDE with previously characterized heterologous DGC and PDE (WspR; RocR), respectively, fully restored their swimming ability to the wild-type levels (Fig. 1B). These results indicate that the presence of putrescine signal exerts a positive effect in regulation of *D. oryzea* swimming motility, whereas the second messenger c-di-GMP plays a negative role in modulation of the bacterial motility.

**FIGURE 1.**
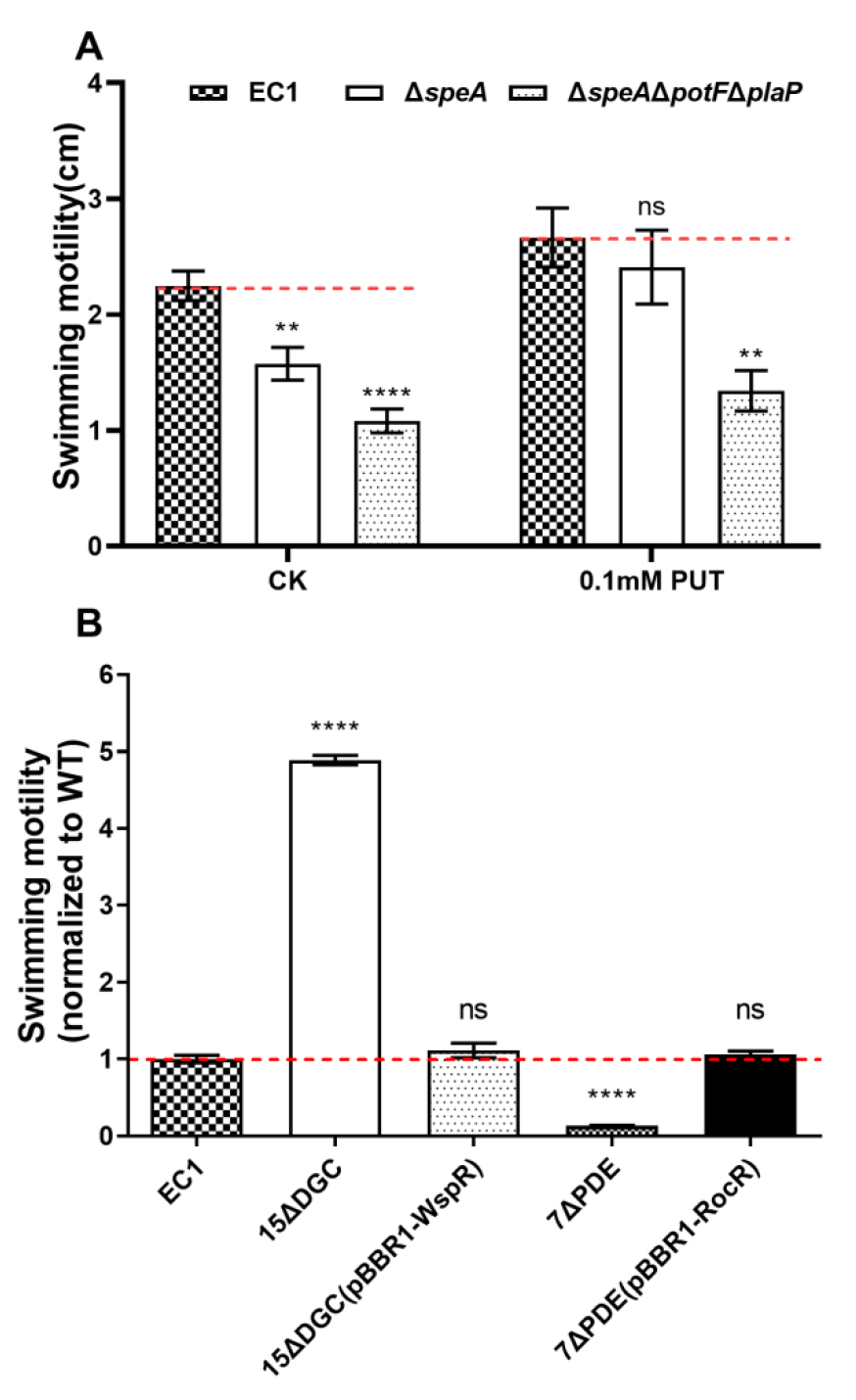
Both c-di-GMP and putrescine can regulate the swimming motility of *Dickeya oryzea* EC1. (A) Putrescine positively regulates the swimming motility of EC1. Δ*speA*, deletion mutant of *speA* gene, encoding arginine decarboxylase; Δ*speA*Δ*potF*Δ*plaP*, the *potF*-*plaP* double disrupted mutant in the *speA*-disrupted genetic background. *potF*, encoding putrescine-binding protein of polyamine ABC transport system potFGHI; *plaP*, encoding a putrescine-specific transporter protein. The final concentration of exogenously added putrescine was 0.1 mM. (B) Negative regulation of swimming motility by c-di-GMP. Δ15DGC, deletion mutant of all c-di-GMP synthase genes in EC1; Δ7PDE, deletion mutant of all c-di-GMP degradation enzyme genes in EC1; Δ15DGC (pBBR1-WspR), complementation of the mutant with a heterologously expressed DGC gene (*wspR*) from *Pseudomonas aeruginosa*; Δ7PDE(pBBR1-RocR), complementation of the mutant with a heterologously expressed PDE gene (*rocR*) from *P. aeruginosa*. Dotted lines indicate the swimming motility of wild-type levels. Experiments were repeated three times in triplicates. ****, *P* < 0.0001; **, *P* < 0.01; ns, *P* > 0.05(by one-way ANOVA with multiple comparisons).

### C-di-GMP accumulation in *D. oryzea* decreased the cellular level of putrescine

In order to discover the potential link between the putrescine communication system and the c-di-GMP signaling pathway, the cellular levels of putrescine and c-di-GMP were measured in the wild type strain EC1 and its corresponding mutants. Intracellular putrescine and c-di-GMP contents were determined by liquid chromatography-mass spectrometry (LC-MS) assays. Results showed that in minimal medium there was no significant change in the c-di-GMP level in the mutant Δ*speA*Δ*potF*Δ*plaP* compared to wild type EC1 (Fig. 2A), suggesting that c-di-GMP metabolism is not influenced by the putrescine signaling system. In contrast, while in LB medium the putrescine levels of the c-di-GMP abundant mutant 7ΔPDE, or its receptor mutant Δ*ycgR,* or double mutant 7ΔPDEΔ*ycgR* and wild type EC1 were comparable at low bacterial cell density (OD_600_ = 0.5 and 1.0), putrescine concentration was dropped significantly by over 50 %, 60 % and 40 % in the mutants Δ*ycgR,* 7ΔPDE and 7ΔPDEΔ*ycgR,* respectively, compared with wild type EC1 along with bacterial growth reaching a high population density (OD_600_ = 1.5) (Fig. 2B). However, there was no significant change in the cellular putrescine content in mutant 15ΔDGC compared to the wild type EC1 under all the bacterial population densities tested (Fig. 2B). The production of putrescine in the complemented strains Δ*ycgR*(*ycgR*) and 7ΔPDE(*RocR*) were recovered to the level of the wild-type strain under the OD_600_ = 1.5 (Fig. 2D). The result of this phenomenon was also present in the minimal medium (Fig. 2C and 2D).

**FIGURE 2.**
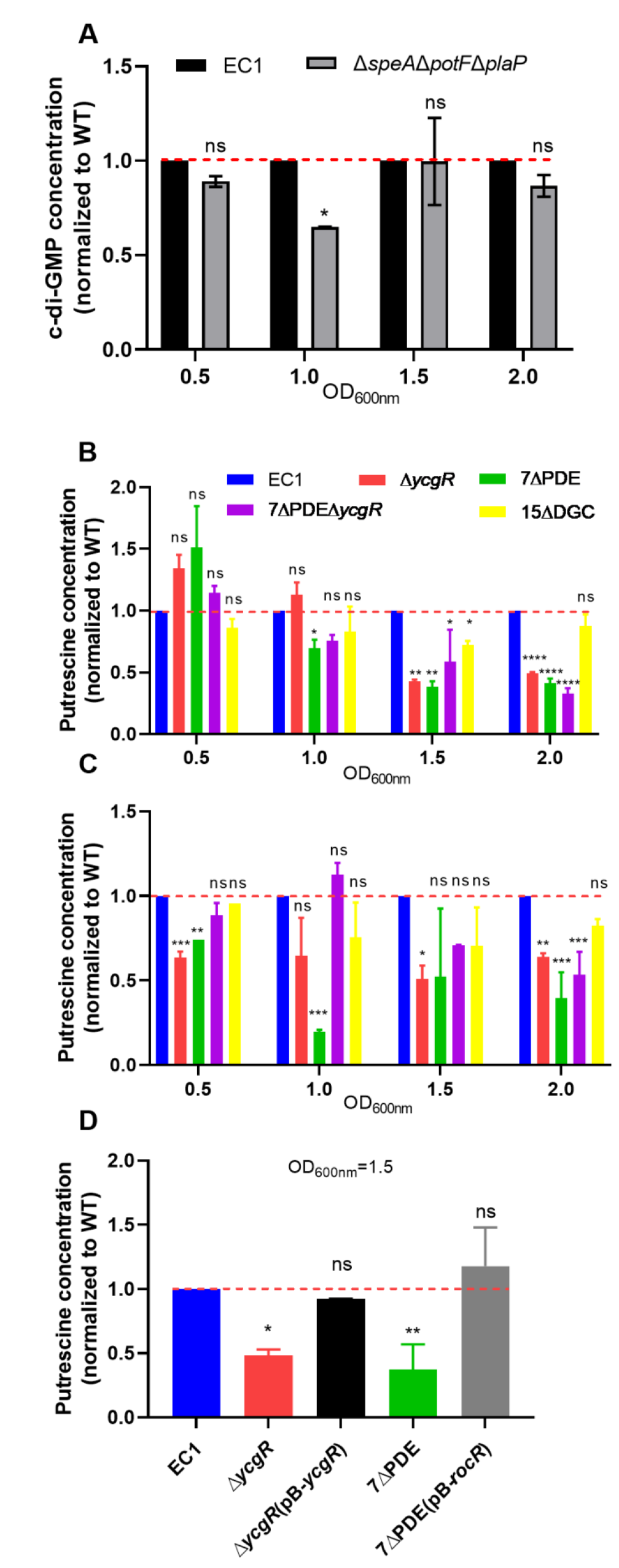
Cellular c-di-GMP and putrescine levels in *Dickeya oryzea* EC1 and its derivatives. (A) Quantitative measurement of cellular cyclic di-GMP concentration of wild-type EC1 and mutant Δ*speA*Δ*potF*Δ*plaP* in Minimal medium. The following genes were quantified in this experiment: *speA*, encoding arginine decarboxylase; *potF*, encoding putrescine-binding protein of polyamine ABC transport system potFGHI; *plaP*, encoding a Putrescine-specific transporter protein. Dotted lines indicate the wild-type levels of c-di-GMP concentrations. *, *P <* 0.05; ns, *P* > 0.05(by Student’s unpaired *t* test). (B) Quantitative measurement of cellular putrescine concentration of strain EC1 and the c-di-GMP signal system-related mutant in LB medium. The following genes were quantified in this experiment: *ycgR*, encoding receptor protein for c-di-GMP; 7PDE, encoding phosphodiesterase that degrade c-di-GMP; 15DGC, encoding diguanylate cyclase that synthesize c-di-GMP; Δ7PDE(pBBR1-RocR), complementation of the mutant with a heterologously expressed PDE gene (RocR) from *P. aeruginosa*; Δ*ycgR*(pBBR1-YcgR); complementation of the mutant with YcgR. ****, *P* < 0.0001; **, *P* < 0.01; *, *P* < 0.05; ns, *P* > 0.05 (by one-way ANOVA with multiple comparisons). (C) Quantitative measurement of cellular putrescine concentration of strain EC1 and the c-di-GMP signal system-related mutant in Minimal medium. Dotted lines indicate the wild-type levels of putrescine concentrations. (D) Quantitative measurement of cellular putrescine concentration of strain EC1, Δ*ycgR*, Δ*ycgR*(pB*-ycgR*)*, 7*ΔPDE, 7ΔPDE(pB-*rocR*). ****, *P* < 0.0001; ***, *P* < 0.001; **, *P* < 0.01; *, *P* < 0.05; ns, *P* > 0.05 (by one-way ANOVA with multiple comparisons). Experiments were repeated three times in triplicates.

To validate the above observations, the transcript levels of the relevant genes encoding the putrescine and ci-di-GMP signaling systems were determined by quantitative reverse transcription-PCR (qRT-PCR) analysis. Given that only three c-di-GMP metabolic genes, i.e., *W909_14945*, *W909_14950* and *W909_10355*, which encode c-di-GMP synthase DdgcA, degradation enzymes DpdeA and DpdeB, respectively, account primarily for the dynamic changes of c-di-GMP content in *D. oryzea* as well as the motility phenotype changes (11), the transcriptional expressions of these genes and the c-di-GMP receptor gene *ycgR* were assessed in wild type strain EC1 and mutant Δ*speA*. The results showed that the expression levels of these genes in mutant Δ*speA* did not increase more than 2-fold and did not decrease more than 50% compared to the wild type level (Fig. 3A). Similarly, the expression levels of the five genes closely related to the putrescine metabolism and transport pathway were also not significantly changed in Δ*ycgR* compared to wild-type strain EC1 (Fig. 3B). According to the above results, c-di-GMP could negatively regulate the production of putrescine, whereas putrescine does not influence the production of c-di-GMP, and the two systems do not affect each other at the transcriptional level.

**FIGURE 3.**
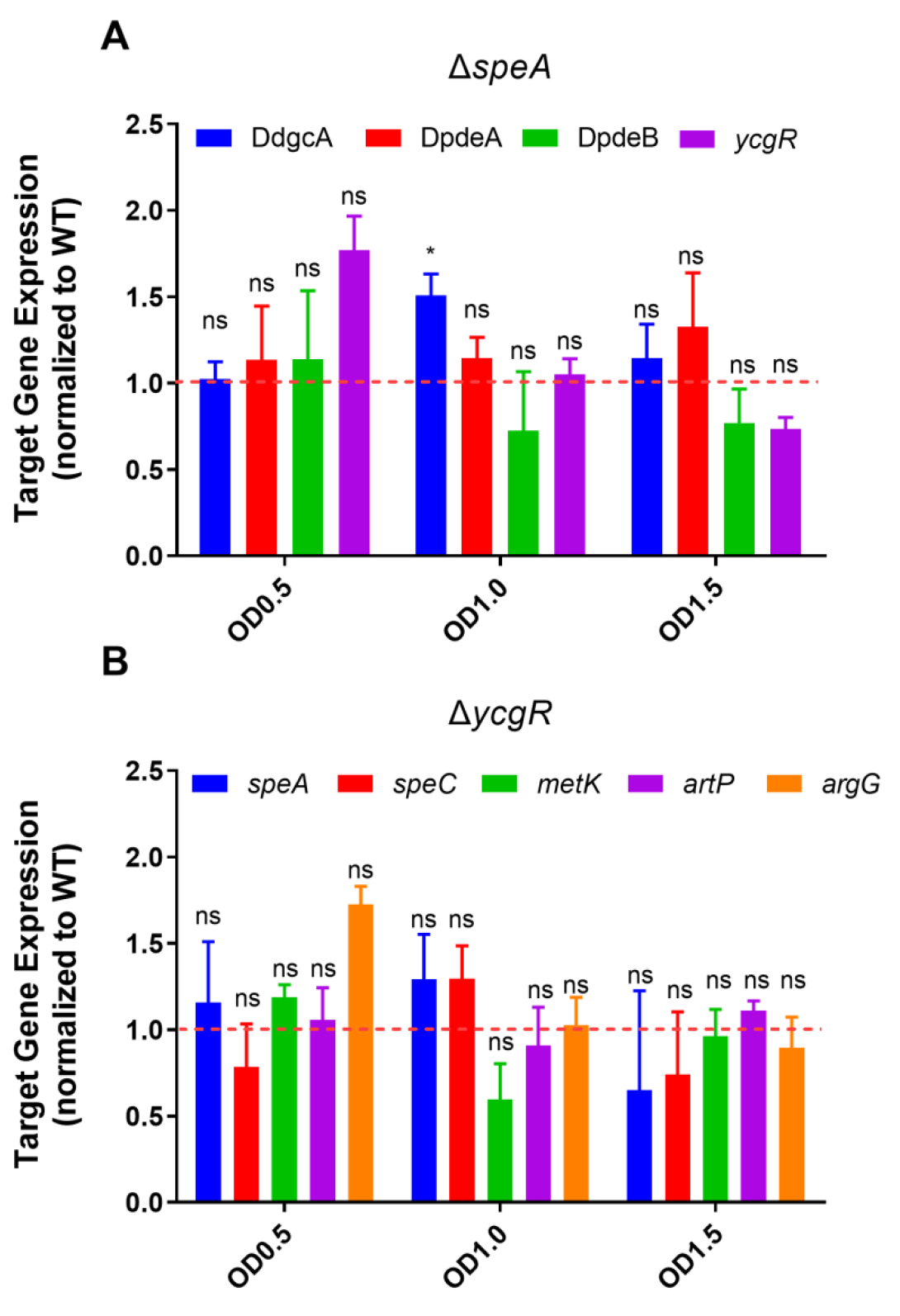
Quantitative reverse transcription-PCR (qRT-PCR) analysis of wild-type EC1 and deletion mutants of two communication systems. (A) Quantification of the expression of *ycgR* gene and three genes encoding the major c-di-GMP synthesis and degradation enzymes in wild-type EC1 and mutant Δ*speA* by qRT-PCR. The following genes were quantified in this experiment: *ycgR*, encoding a PilZ domain receptor for c-di-GMP; DdgcA(W909_14945), encoding the major diguanylate cyclase that synthesize c-di-GMP; DpdeA(W909_14950), encoding the major phosphodiesterase that degrade c-di-GMP; DpdeB(W909_10355), encoding the major bifunctional enzymes that can both synthesize and degrade c-di-GMP(9). (B) Quantification of the expression of the putrescine pathway-related genes in wild-type EC1 and mutant Δ*ycgR* by qRT-PCR. *speA* and *speC* encode two key rate-limiting enzymes for putrescine synthesis, proteins encoding by speA and other three genes could interact with YcgR. The following genes were quantified in this experiment: *speA*, encoding arginine decarboxylase; *speC*, encoding ornithine decarboxylase; *metK*, encoding S-adenosylmethionine synthase; *artP*, encoding arginine ABC transporter ATP-binding protein; *argG*, encoding argininosuccinate synthase[]. The dotted lines represent the gene expression levels in wild type, which was set to a value of 1. *, *P* < 0.05; ns, *P* > 0.05(by multiple *t* test) (n ≥3 independent experiments).

### Interaction between YcgR and putrescine metabolic proteins

From results in the previous section, it is clear that c-di-GMP can regulate the levels of putrescine negatively (Fig. 2). In general, c-di-GMP molecules are dependent on downstream receptors to modulate the corresponding phenotype. And YcgR class receptor was mainly responsible for modulation of bacterial motility in EC1 via its PilZ structural domain(42). Considering the above, this section focused on the interaction of the c-di-GMP receptor YcgR with putrescine pathway proteins.

Using the immunoprecipitation-Mass Spectrometry (IP-MS) and bioinformatic analysis, seven proteins related to putrescine metabolic pathway that interacted with YcgR were identified (Table 1), among them, four proteins were further validated by bacterial two-hybrid assay (B2H), namely, SpeA, ArgG, ArtP and MetK (Fig. 4A). SpeA is able to catalyze the synthesis of agmatine from L-arginine(8). ArgG can catalyze the synthesis of L-arginino-succinate acid from citrulline(43). ArtP is a transporter protein for the polyamine synthesis precursor L-arginine and is capable of transferring extracellular L-arginine into the cell(44). MetK is able to catalyze the synthesis of S-adenosylmethionine from L-methionine(45), which can further react with putrescine to form spermidine. From the result of IP-MS, the strongest interaction was between MetK and YcgR, followed by SpeA and ArgG (Table 1). Surprisingly, the result of B2H showed interaction between MetK and YcgR was not the strongest as IP-MS indicated (Fig. 4A). Swimming motility of deletion mutants of these four proteins showed that only Δ*speA* was significantly less motile compared to the wild type, while Δ*argG*, Δ*artP* and Δ*metK* was comparable with the wild type (Fig. S1). According to the results above, there are four proteins involved in the putrescine metabolic pathway that can interact with YcgR.

**FIGURE 4.**
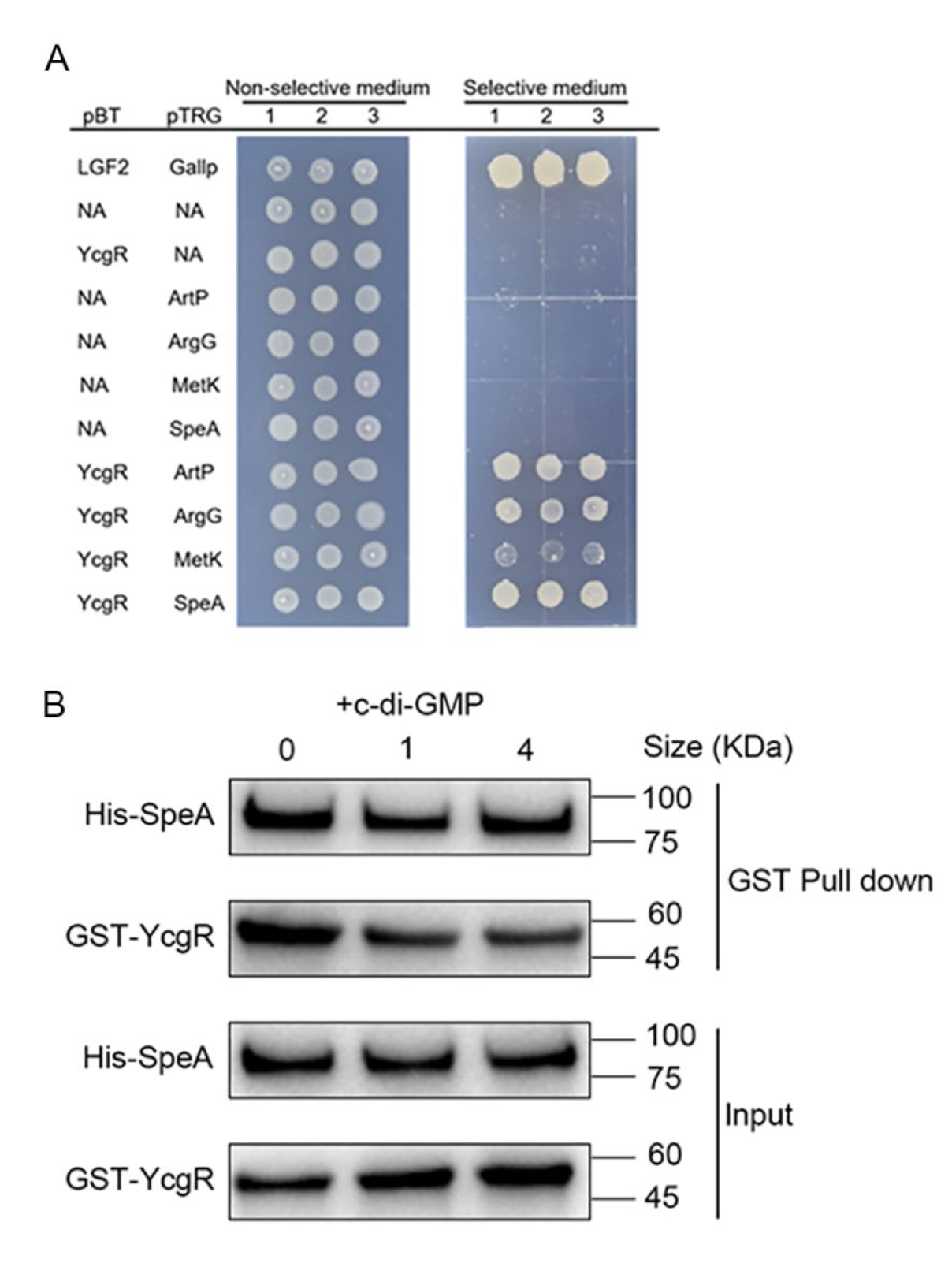
YcgR is able to interact with four putrescine pathway-related proteins. (A) Putrescine pathway-associated proteins capable of interacting with YcgR were further validated by bacterial two-hybridization. Four putrescine pathway-related proteins that can directly interact with YcgR have been identified in this experiment as follows: SpeA, biosynthetic arginine decarboxylase; ArtP, arginine ABC transporter ATP-binding protein; ArgG, argininosuccinate synthase; MetK, S-adenosylmethionine synthase. (B) GST pull-down experiment analysis of interaction between YcgR and SpeA. M: marker; lane 1: purified YcgR; lane 2: purified SpeA; lane 3: purified YcgR:SpeA interaction.

**Table 1.** Partial proteins that interact with YcgR and are associated with the metabolic pathway of polyamines were identified by Co-IP and protein profile identification.

| GeneName | ORF names | ProteinName | UniquePepCount |
| --- | --- | --- | --- |
| argG | W909_00315 | Argininosuccinate synthase, L-arginine biosynthesis | 3 |
| argA | W909_04780 | Amino-acid acetyltransferase | 1 |
| artP | W909_05770 | Arginine ABC transporter ATP-binding protein | 1 |
| plaP | W909_08760 | Putrescine/spermidine ABC transporter | 1 |
| strP | W909_10855 | Polyamine ABC transporter permease | 1 |
| speA | W909_17465 | Biosynthetic arginine decarboxylase | 4 |
| metK | W909_17475 | S-adenosylmethionine synthase | 9 |

### YcgR can enhance the production of putrescine by promoting the activity of SpeA *in vitro*

Considering that SpeA is a key rate-limiting enzyme for putrescine synthesis, and only Δ*speA* mutant decreased swimming motility significantly in EC1 (Fig. S1) this section therefore focused on the interaction mechanism between YcgR and SpeA. In *D. oryzea* EC1, SpeA can catalyze the conversion of arginine to agmatine, which in turn is converted into putrescine in a two-step reaction(8). In this study, the concentration of agmatine was measured using OPA (*o*-phthalaldehyde) spectrofluorometric method (46, 47). Results showed that the addition of YcgR to the reaction system can increase the production of agmatine compared to when only SpeA was added, and when YcgR was added to the reaction system at a molar ratio of 1:2 between SpeA and YcgR, the increase in production of agmatine was the most significant (Fig. 5A and Fig. S5). In detail, when YcgR was added to the reaction for 1 h, the yield of agmatine increased by 75% compared to the null-YcgR group; after the reaction was carried out for 2 h, the agmatine production of the YcgR-added group greatly increased by almost 1.7 folds (Fig. 5A). It suggested that this promotion effect was more obvious with the increase of reaction time. In previous studies, we have shown that YcgR binds c-di-GMP in a 1:1 molar ratio (9), therefore, the effect of c-di-GMP to the interaction between SpeA and YcgR was also investigated. Results showed that the addition of c-di-GMP to the enzymatic reaction resulted in a significant decrease in the production of agmatine compared to the group without adding c-di-GMP (Fig. 5B). From the above results, it appears that YcgR can promote the enzyme activity of SpeA by increasing the production of its product agmatine, whereas c-di-GMP appears to have an inhibitory effect on this promoting effect.

**FIGURE 5.**
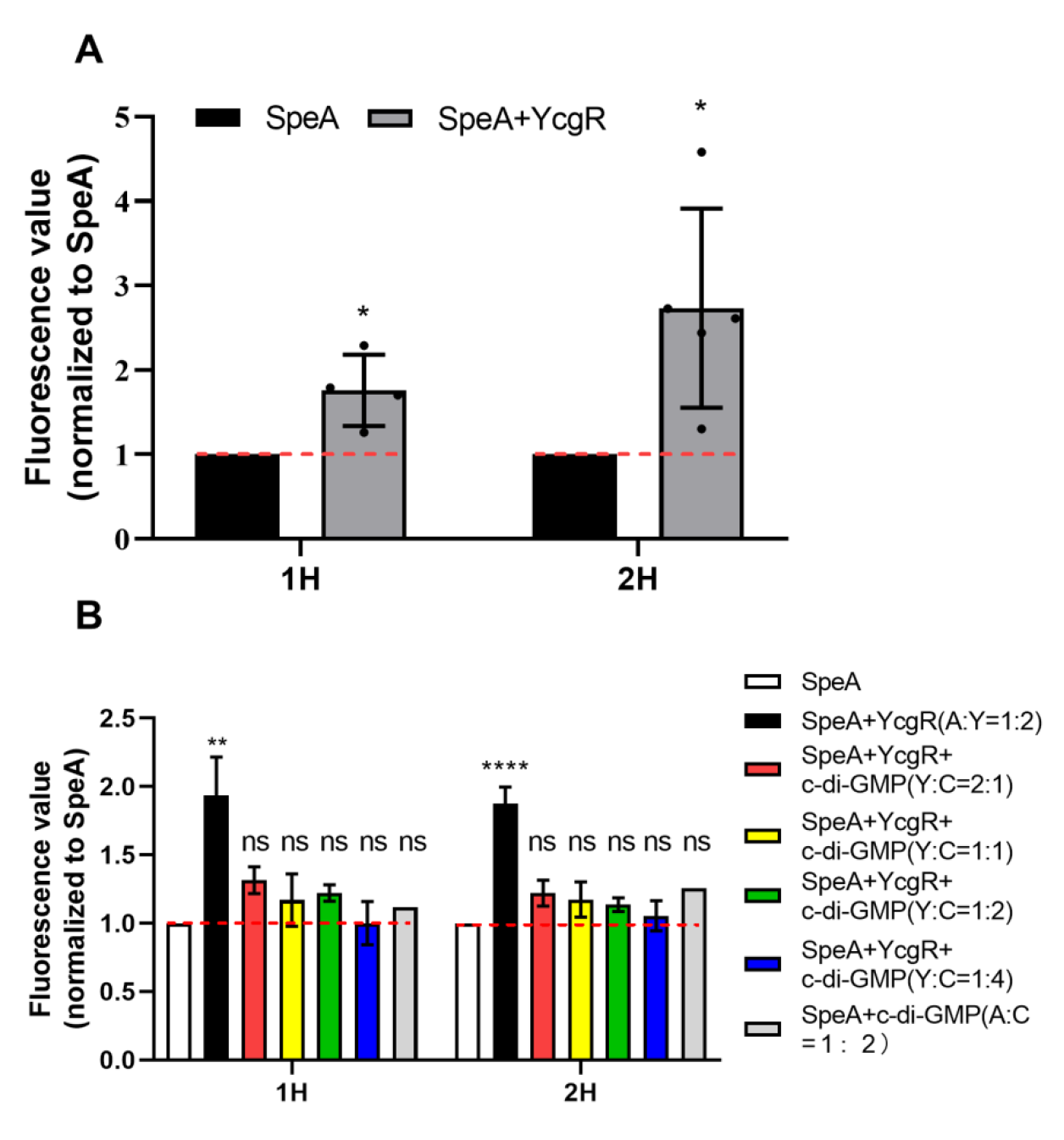
*In vitro* YcgR interacts with SpeA and promotes its activity as an arginine decarboxylase(ADC). (A) Enzyme activity assay of SpeA *in vitro* indicated that YcgR promoted the ADC activity of SpeA. The product agmatine was extracted after 1h and 2h respectively of the reaction and its absorbance value was measured after 8h. Dotted lines indicate the agmatine production of the system when only SpeA was added. *, *P* < 0.05; ns, *P* > 0.05(by Student’s unpaired *t* test). (B) Binding to c-di-GMP inhibited the promotional effect of YcgR on the ADC activity of SpeA. Dotted lines indicate the agmatine production of the system when only SpeA was added.****, *P* < 0.0001; **, *P* < 0.01; ns, *P* > 0.05(by one-way ANOVA with multiple comparisons). All experiments were repeated three times in triplicates.

### C-di-GMP plays a major role in regulating swimming motility in comparison with putrescine

It is clear that both the putrescine communication system and the second messenger c-di-GMP can regulate the bacterial motility in EC1, so this section focused on which signal system dominates in the regulation of the bacterial motility. Firstly, a heterologous PDE RocR from *P. aeruginosa* (48) was overexpressed in Δ*speA*Δ*potF*Δ*plaP* to observe whether the bacteria could restore its swimming motility in the complete absence of putrescine background, which reversed the motility of Δ*speA*Δ*potF*Δ*plaP* to the wild type level (Fig. 6A). On the other hand, after adding putrescine at a final concentration of 0.1 mM, only the swimming motility of Δ*speA* could restore to wild-type level, while 7ΔPDE remained nonmotile (Fig. 6B). And the deletion of the *speA* gene in 15ΔDGC also did not reduce its already increasing motility (Fig. 6C).

**FIGURE 6.**
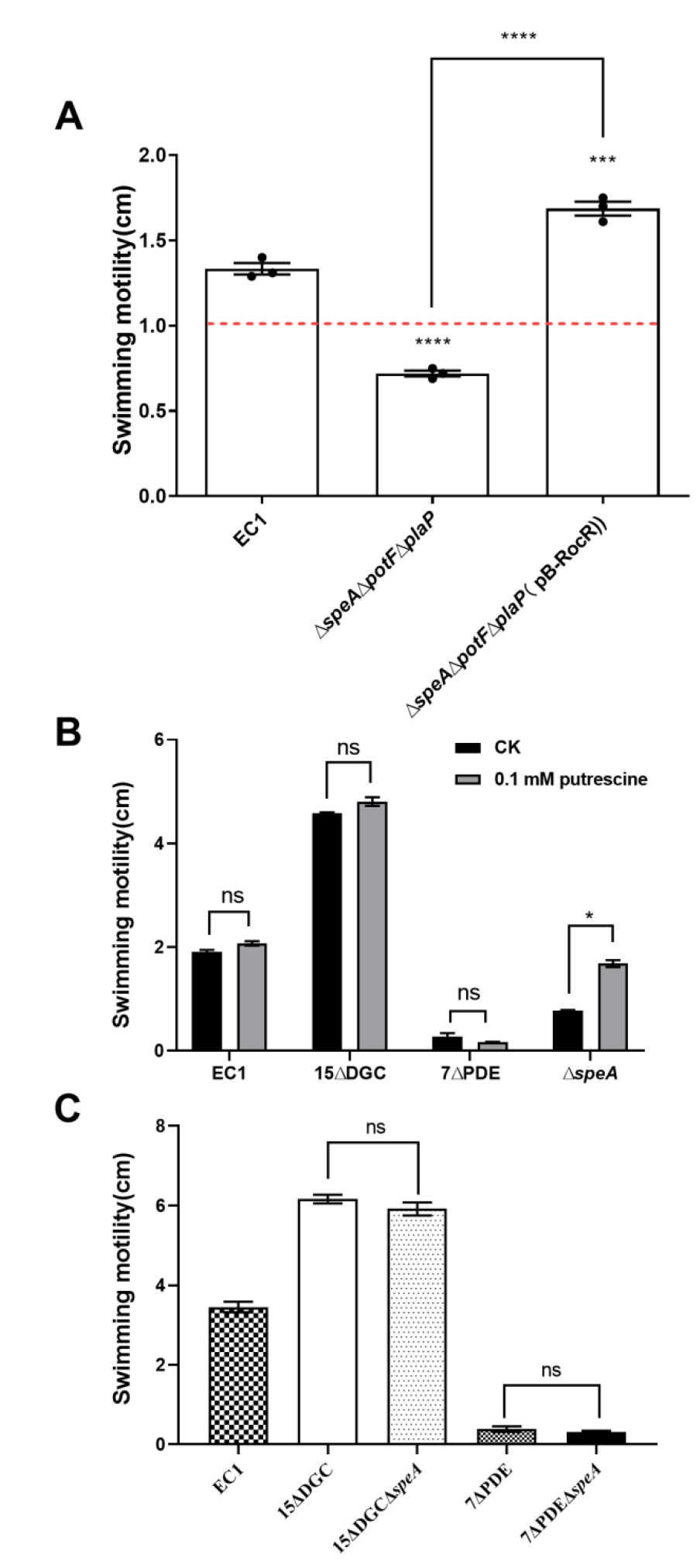
The dominance of c-di-GMP over putrescine in the regulation of bacterial motility. (A) Overexpression of c-di-GMP degradation enzymes restored the swimming motility of Δ*speA*Δ*potF*Δ*plaP*(Δ*3A*). RocR: c-di-GMP phosphodieasterase of *P. aeruginosa*. Dotted lines indicate the diameter of swimming zone of wild-type EC1. Experiments were repeated three times in triplicates. ****, *P* < 0.0001; ***, *P* < 0.001 (by one-way ANOVA with multiple comparisons). (B)Swimming motility of strain EC1,15ΔDGC,7ΔPDE and Δ*speA*. The final concentration of exogenously added putrescine was 0.1 mM. Experiments were repeated three times in triplicates. *, *P* < 0.05; ns, *P* > 0.05(by Student’s unpaired *t* test). (C) Swimming motility of strain EC1,15ΔDGC,15ΔDGCΔ*speA*,7ΔPDE and 7ΔPDEΔ*speA*. Experiments were repeated three times in triplicates. ns, *P* > 0.05(by one-way ANOVA with multiple comparisons).

In summary, c-di-GMP signaling pathway occupies a dominant position in the regulation of bacterial motility compared to the putrescine communication system.

## Discussion

In this study, we investigated the interaction between two signal systems, the putrescine communication system and the c-di-GMP signal system. Both of these two signaling systems regulate bacterial motility of EC1 (Fig. 1). The levels of c-di-GMP molecules in Δ*speA*Δ*potF*Δ*plaP* were not significantly different compared to the wild type EC1 (Fig. 2A). However, when the cell density reached mid-log phase, the concentration of putrescine was decreased significantly in Δ*ycgR,* 7ΔPDE and 7ΔPDEΔ*ycgR* respectively compared with wild type level (Fig. 2B and 2C), which suggested that both the deletion of the c-di-GMP receptor protein YcgR and high levels of c-di-GMP could decrease putrescine concentration. Then, multiple experimental approaches such as, IP-MS, bacterial two-hybrid and pull-down assay, were used to identified an interaction between YcgR and SpeA (Table 1 and Fig. 4). *In vitro* enzymatic assay showed that YcgR could enhance the production of agmatine by promoting the ADC activity of SpeA, which could be inhibit by adding c-di-GMP molecule into the reaction system (Fig. 5). In addition, our results suggest c-di-GMP plays a major role in regulating swimming motility in comparison with putrescine(Fig. 6). Taken together, the data from this study demonstrate that c-di-GMP could interact with putrescine via a PilZ domain receptor YcgR, whcih could enhance the production of putrescine by promoting the activity of SpeA, and this facilitative effect could be inhibited by c-di-GMP molecules (Fig. 7).

**FIGURE 7.**
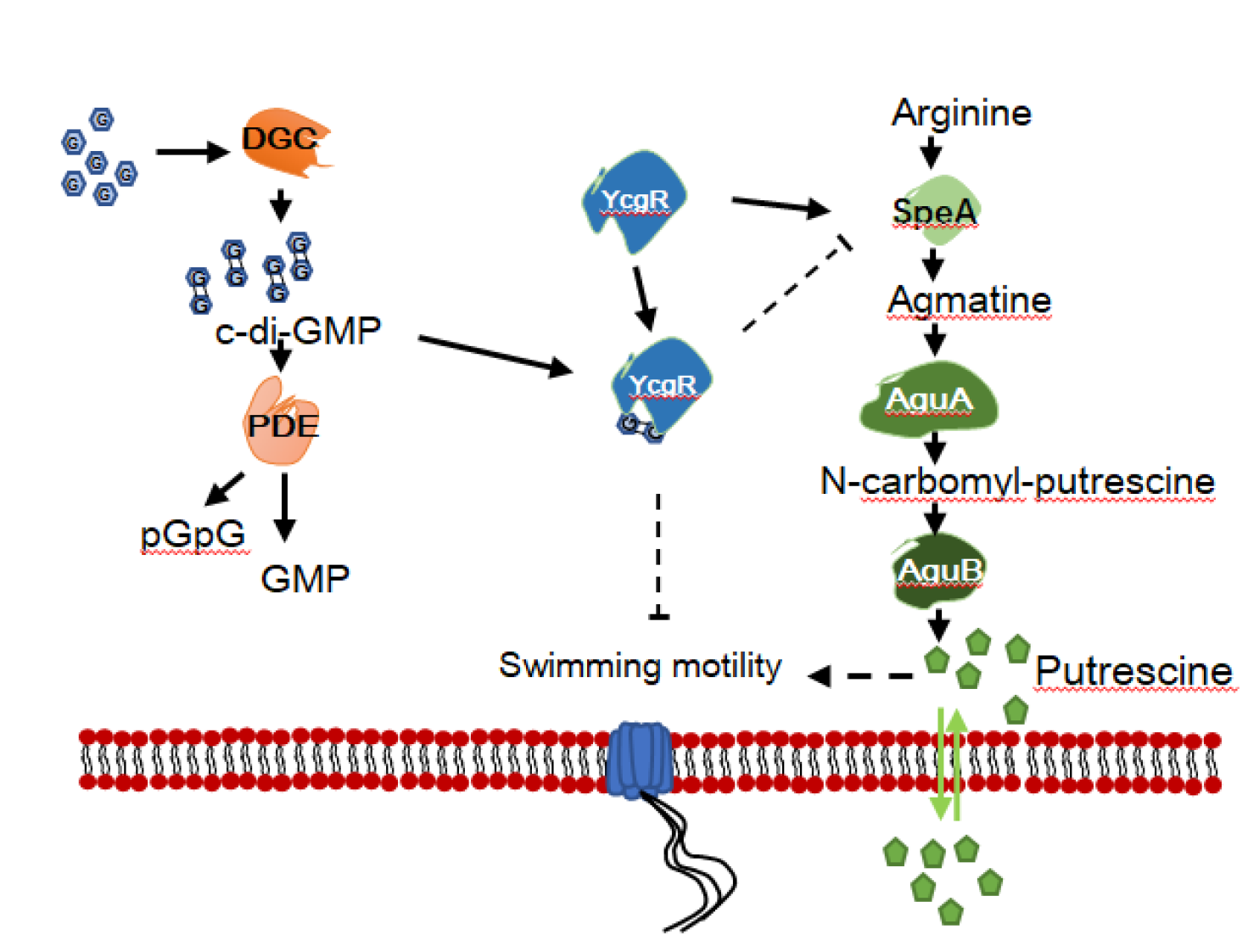
The regulatory model between the putrescine communication system and c-di-GMP signal system in EC1. SpeA, arginine decarboxylase; AguA, agmatine deiminase; AguB, N-carbamoylputrescine aminotransferase; YcgR, a PilZ domain receptor for c-di-GMP.

This regulatory model in EC1 is different from previous studies in other bacteria. In *P. aeruginosa*, putrescine can induce the accumulation of intracellular c-di-GMP and promote biofilm formation(49). In *Agrobacterium tumefaciens*, where polyamines control the formation of biofilm by regulating the concentration of c-di-GMP(50). In *Vibrio cholerae*, a recent study presented a model in which norspermidine and spermidine could competitively bind the periplasmic protein NspS, thereby allowing NspS to regulate the bifunctional activity of MbaA, which synthesizes c-di-GMP and degrade c-di-GMP in the presence of norspermidine and spermidine, respectively(51). Bioinformatic analysis showed that there was no gene homologous of YcgR in *P. aeruginosa, A. tumefaciens* and *V. cholerae,* and EC1 did not contained homologs of MbaA from *V. cholerae*, which may partly explain why the regulatory model in EC1 differs from these bacteria. Another possible reason is that EC1 is evolutionarily distant from these bacteria, and *P. aeruginosa* and *V. cholerae* are important pathogen for humans and animals, whereas strain EC1 is a plant pathogenic bacterium. Furthermore, whether the two communication networks are mutually regulated at the transcriptional level was further investigated by qRT-PCR, which showed no transcriptional interaction between these two systems (Fig. 3). These features suggest the complexity and specificity of the interaction between putrescine and c-di-GMP in different bacterial species, highlighting the necessity to elucidate this interaction.

How does c-di-GMP affect the putrescine communication system in EC1? In screening for proteins that can interact with YcgR we found that it was able to interact with four putrescine biosynthesis pathway-related proteins, especially SpeA, the rate-limiting enzyme of the putrescine biosynthesis pathway (Fig. 4). *In vitro* experiments showed that YcgR could promote the ADC activity of SpeA by enhancing the production of agmatine (Fig. 5A). The increase in agmatine production will inevitably result in a increase in intracellular putrescine levels(52). This result is consistent with the observation that Δ*ycgR* mutant significantly decrease putrescine (Fig. 2). The decrease in putrescine production in 7ΔPDE is another question to consider, that is in what way does the large accumulation of c-di-GMP contribute to the decrease in putrescine production. The addition of different concentrations of c-di-GMP molecules to the enzyme reaction system of SpeA and YcgR resulted in a decrease in agmatine production with increasing final concentration of c-di-GMP(Fig. 5B). Given that YcgR is able to bind to c-di-GMP molecules in a molar ratio of 1:1(9), it may be possible that was due to the high concentration of c-di-GMP preferentially binding to YcgR and thus prevents it from binding to SpeA, thereby inhibiting the facilitative effect of YcgR on SpeA enzyme activity. In previous reports, YcgR-like pilz protein was found to regulate motility mainly through direct interaction with the bacterial flagellar motor proteins FliM, FliG and MotC, etc, or through protein-DNA interactions at the transcriptional level affecting the expression of genes involved in bacterial biofilm formation to regulate the phenotype of the bacterial attachment state(27, 53, 54). These results led to the first knowledge of that YcgR-like protein acts as a receptor for c-di-GMP to regulate motility by interacting with putrescine biosynthesis pathway-related proteins. All of the above features demonstrate the importance of YcgR in linking the two signal systems in regulating motility. However, identification of the specific interplay domains between YcgR and SpeA remains to be further investigated. In-depth study and understanding of the structural domains of the two proteins and their interaction mechanisms is also important to explain the most significant facilitation effect of YcgR and SpeA at a molar ratio of 2:1.

This study also determined that c-di-GMP dominates in the regulation of bacterial motility over putrescine in EC1. The importance of putrescine and c-di-GMP in the regulation of bacterial motility has been highlighted in numerous previous studies(8, 9, 53, 55, 56), but there is no report on which signaling system is more dominant in terms of regulating motility. Our results showed that the motility of both Δ*speA*Δ*potF*Δ*plaP* and 7ΔPDE was significantly reduced compared to the wild type EC1, while the motility of 15ΔDGC was significantly enhanced compared to EC1. And *in trans* expression of the c-di-GMP degrading enzyme in Δ*speA*Δ*potF*Δ*plaP* was able to restore its motility to above wild-type level, whereas adding 0.1 mM of putrescine did not restore the motility of 7ΔPDE and the deletion of the *speA* gene in 15ΔDGC did not reduce its motility(Fig. 6), indicating that c-di-GMP dominates putrescine in the regulation of bacterial motility. The pattern of YcgR as a braking protein to regulate motility is conserved in most bacteria, and it is mainly achieved through the interactions between YcgR and flagellar structural proteins such as FilG and MotA(57). However, we found that in EC1, in addition to regulating motility by interacting with SpeA, YcgR could also interact with bacterial chemotaxis proteins, but not with flagellar structural proteins(Fig. S6). Chemotaxis proteins have been reported to control the direction of bacterial flagellar rotation by directly interacting with the flagellar protein FliG, FliM, and FliN(58); it indicates the importance of chemotactic proteins in the regulation of bacterial motility. We therefore speculate that YcgR may primarily regulate bacterial motility by interacting with chemotactic proteins, but this possibility still needs to be further tested.

From the previous research, in addition to putrescine and c-di-GMP, the bacterial motility of *D. oryzea* EC1 also regulated by the AHL quorum sensing system and Vfm-mediated quorum sensing system(32, 59), suggests the complexity of motility regulation in EC1. As for the regulatory relationship between these singnalling system, it is still under further investigate to clarify. Whether these signalling systems are linked together and thus collectively regulate motility or individually, this will all contribute to the refinement of the bacterial motility regulatory network and provide new insights and references for subsequent studies.

Taken together, these findings shed light on the relationship between putrescine and c-di-GMP signaling network. Both of them could regulate the bacterial motility of EC1, and disruption of the c-di-GMP signalling system significantly reduced the production of putrescine in bacteria. We showed that c-di-GMP interact with putrescine via a PilZ domain receptor YcgR. It has been demonstrated that YcgR can facilitate putrescine production by promoting SpeA’s enzymatic activity, but c-di-GMP molecules can inhibit this facilitative effect. Furthermore, we also indicated the dominance of c-di-GMP over putrescine in the regulation of bacterial motility. To our knowledge, this is the first time that the interaction between c-di-GMP and the putrescine communication system has been revealed and clarified. Additionally, these features have also enhanced our understanding of downstream regulatory modes and pathways of YcgR-like proteins.

## MATERIALS AND METHODS

### Bacterial strains and plasmids

Bacterial strains and plasmids used in this research are listed in Table S1 in the supplemental materials. *Escherichia coli* was routinely grown at 37°C in Luria-Bertani (LB) medium. *D. oryzea* EC1 and its derivatives were grown at 28°C in LB medium as previously reported. Minimal medium (MM) agar plates were used for conjugation(32). The following antibiotics were added at the indicated final concentrations when required: ampicillin (Ap) at 100 µg/ml, kanamycin(Km) at 50 µg/ml, streptomycin (Str) at 50 µg/ml, and polymyxin (Pm) at 30 µg/ml. The optical density at 600 nm (OD_600_) of the bacterial culture was measured by using a NanoDrop 2000c system (Thermo Fisher Scientific, USA) at 600 nm.

### Mutant construction and complementation

The generation of in-frame gene deletion mutants was performed using the suicide vector pKNG101 and triparental mating according to a previously described protocol(60). In-frame deletion of the coding regions of all genes were done by the allelic-exchange method(61). Flanking regions of each coding region were amplified by PCR using the specific primers listed in the Table S2 in the supplemental materials. A complementation assay was performed by using the plasmid pBBR1MCS-4 and triparental mating, according to a previously described protocol(60). The coding region of the target gene was amplified by PCR using the specific primers (Table S2).

### Bacterial motility assay

Collective swimming motility was assessed in a semisolid medium plate with 0.2% agar (each liter containing 10 g Tryptone, 5 g yeast extract, 10 g NaCl and 2 g agar). A bacterial culture grown to an OD_600_ of 1.0 (1 µl) was spotted on the center of the plate and incubated at 28°C for 12 to 18 h before measurement. Collective swarming motility was assayed as previously described(60).

### Quantitative analysis of c-di-GMP by LC-MS

Quantification of c-di-GMP levels in wild type strain EC1 and mutant Δ*speA*Δ*potF*Δ*plaP* were measured based on the methods described previously by Chen without modification(9). Cellular c-di-GMP concentration was determined by perchloric acid lysis and liquid chromatography-mass spectrometry (LC-MS) in wild type EC1 and mutants related to the putrescine communication system. The c-di-GMP samples were separated by using an Syncronis C18 column (Thermo Fisher Scientific, USA) fitted with a 100- by 2.1 mm guard column with a flow rate of 0.2 ml/min, and the cycle time was 10 min. c-di-GMP was detected with an orbitrap mass analyzer on the Q Exactive Focus system (Thermo Fisher Scientific, USA) in positive ionization mode. c-di-GMP levels were normalized to the total protein per milliliter of culture. Data represent the means from three independent cultures, with error bars indicating the standard deviations. The experiments were repeated three times in triplicates.

### Quantitative analysis of putrescine by LC-MS

Quantification of putrescine levels in wild type strain EC1 and mutants of c-di-GMP systems were measured based on the methods described previously with minor modifications (62–65). Cells were grown overnight in LB medium at 28°C, adjusted to an OD_600_ of 1.0, and then subcultured in 15-ml minimal medium with a 100-fold dilution in a 50-ml culture tube (Crystalgen, USA). When the bacterial culture was grown to an OD_600_ of target value (0.5, 1.0, 1.5 and 2.0), an aliquot of 2 ml was transferred into a 15-ml centrifuge tube. The bacteria were then collected by centrifugation at 4,000 rpm for 5 min and 2 ml xTractor Buffer (Clontech, TaKaRa Biomedical Technology [Beijing] Co., Ltd., China) was added to each tube to lyse the bacteria. 800 µl of the lysed sample was then taken in a new 15-ml centrifuge tube. For each sample, derivatization was carried out by adding 1 ml of 2 M NaOH and 100 µL of benzoyl chloride and vortexing 2 min and incubated for 20 min at 37 ℃. 2ml of saturated sodium chloride solution was added to each tube and vortexed for 2min, followed by adding 2ml of petroleum ether for extraction. The samples were extracted overnight at 4℃ and then centrifuged at 4,000 rpm for 5 min at 4°C using a 5810 R fixed-angle rotor centrifuge (Eppendorf, Germany). 1 ml of the upper layer of petroleum ether was dried under vacuum in a new 2-ml tube, followed by the addition of 500 µl of methanol to fix the volume. After filtering the organic solvent through a 0.22 µm filter, all samples were stored at −20℃. Benzoylated putrescine was used for analysis with high performance liquid chromatography mass spectrometry (LC-MS). The derivatized samples were separated by using an ACQUITY UPLC HSS T3 column (Waters, Milford, MA, USA) fitted with a 100- by 2.1 mm guard column with a flow rate of 0.3 ml/min. Q Exactive Focus system (Thermo Fisher Scientific) was used to verify the identity of each peak observed in HPLC fractions. The experiments were repeated three times in triplicates. A standard curve was generated by using various concentrations of benzoylated putrescine in duplicates.

### Protein expression and purification

The coding sequence of *speA* was amplifified by PCR using EC1 genomic DNA as the template and then cloned into the pET-32a(+) expression plasmid at the BamHI and HindIII sites using a ClonExpress II one-step cloning kit (Vazyme Biotech Co., Ltd., China). And the coding sequence of *ycgR* was amplifified by PCR using EC1 genomic DNA as the template and then cloned into the Pgex-6P-1(+) expression plasmid at the XhoI and BamHI sites using the same cloning kit. Recombinant plasmids were respectively transformed into *E.coli* DH5α competent cells for sequencing, and the correct constructs of pET-speA and pET-ycgR were respectively transformed into *E.coli* BL21(DE3) for fusion protein expression. For the overexpression of fusion proteins, cultures of expression strains grown overnight were inoculated into 200ml LB medium in 500-ml Fernbach flask and grown with shaking until the OD_600_ reached ∼0.6 to 0.8, isopropyl-β-D-thiogalactopyranoside (IPTG) was added to a final concentration of 0.5 mM to induce the expression of fusion proteins, and cultures were placed in a shaker at 16℃ grown by shaking overnight. Cells were harvested by centrifugation at 4,000 rpm for 20 min at 4℃ and resuspended in 20 ml xTractor buffer (Clontech, TaKaRa Biomedical Technology [Beijing] Co., Ltd., China) for lysis. Cell suspensions were incubated at room temperature for 10 min with gentle shaking. Crude lysates were centrifuged at 4,000 rpm for 90 min at 4°C, and supernatants were filtrated using 0.45- µm syringe filters (Pall Corporation). Affinity purification was performed at 4°C using Talon metal affinity resins (Clontech, TaKaRa Biomedical Technology [Beijing] Co., Ltd., China).

For the fusion protein SpeA, the resin and the column were washed with 10 column volumes of equilibration buffer (50 mM sodium phosphate, 300 mM sodium chloride, 20 mM imidazole [pH 7.4]) before adding the supernatant lysate. The target protein was eluted with gradient elution buffer (50 mM sodium phosphate, 300 mM sodium chloride, 50/100/150/200/300 mM imidazole [pH 7.4]). And for the fusion protein YcgR, the resin and the column were washed with 10 column volumes of equilibration buffer (1ⅹPBS buffer [pH 7.4]) before adding the supernatant lysate. The target protein was eluted with elution buffer (10 mM reduced glutathione, 50 mM Tris-Hcl [pH 7.4]). All the fractions were estimated by Coomassie brilliant blue staining after SDS-PAGE. The protein concentration was determined at 280 nm using the NanoDrop 2000c system (Thermo Fisher Scientific, USA).

### Enzyme activity assay of SpeA in vitro

The ADC enzyme activity of SpeA was measured according a prcedure described previously by song et al.(66), with modifications. The reaction mixture of SpeA activity contained 5.767ⅹ10^-4^µM SpeA protein and 0.1 µg L-arginine in 0.5 ml reaction buffer (50 mM HEPES2.5 mM MgSO_4_、3.3 mM DTT、0.5 mM PLP, PH[8.0]). And YcgR was added in molar ratios of 1:0, 1:1, 1:2, and 2:1, respectively, to SpeA. As for c-di-GMP, it was added in molar ratios of 2:1, 1:1, 1:2, and 1:4, respectively, to YcgR.

The mixture was incubated in a water bath at 37℃ for 2 h, and 200 µl of each sample was taken out every 1 h and quenched by adding 200 µl NaCl-saturated 10% KOH solution, then 200 µl n-butanol were adding for extraction of agmatine. After 8 h extraction, 20 µl of the top n-butanol layer was removed and mixed with 200 µl OPA (o-phthalaldehyde) reagent (0.8mg/ml of o-phthalaldehyde (OPA), [pH 10.0])to detect colored derivatives in this layer. The accumulation of agmatine can be measured the fluorescence at excitation 340 nm and emission at 460 nm on a Bioscreen-C Automated Growth Curves Analysis System.

### Immunoprecipitation-Mass Spectrometry (IP-MS) and bioinformatic analysis

Cell pellets from the EC1 (GFP-YcgR) strain was harvested and washed twice with PBS buffer and resuspended in PBS to a final OD _600nm_ = 4. Add cross-linker DTBP (Thermo Scientific, catalog no. 20665) in the cell suspension to a final concentration of 5 mM and incubated for 45 min at room temperature. Stop cross-linking reaction by adding Tris-HCl to a final concentration of 20 mM for 15 min. Then total lysates from the EC1(GFP-YcgR) strain were extracted by using the French Pressure cell(100Mpa, 1 passage) with 20 U/ml DNaseI and proteinase inhibitor cocktail (Thermo Scientific, catalog no. 87786). Such nondenatured lysates were subject to immunoprecipitation with GFP-Trap (Chromo Tek, gta-20). The IP proteins (about 30 μg) were digested in solution using trypsin for 20 h at 37°C, and the digested peptides were subjected to Nano-HPLC/ESI-ion trap-MS/MS analysis with a Q Exactive mass spectrometer (Thermo Fisher Scientific, US). Raw MS data files were processed and analyzed using Mascot2.2 (Matrix Science, UK) for database search with the following parameters: Database, Dickeya_zeae_NZ_CP006929.1; Enzyme, trypsin; Variable modifications, Oxidation (M); Fixed modifications, Carbamidomethyl (C); Missed Cleavages, 2; Peptide Mass Tolerance, 20 ppm; Fragment Mass Tolerance, 0.1 Da; Filter by score, ≥20.

### The bacterial two-hybrid assay

The bacterial two-hybrid assay to assess the presence of protein-protein interactions between YcgR and putrescine pathway associated proteins was conducted as previous described(67), by using the BacterioMatch II two-hybrid system kit (Stratagene, San Diego, CA). In this assay, approximately 50 μg of each plasmid harboring pBT-*ycgR* and pTRG-*artP/argG/metK*/*speA* were co-transformed into the host strain, XL1 - Blue MRF’ Kan, through chemical transformation after adding 1 mL of pre-heated LB medium in each tube. The cells were then incubated at 37 °C while shaking at 200 rpm for 2 h. Thereafter, the LB medium was removed through centrifugation then the cells were resuspended in 1 mL of M9+ His-dropout broth and incubated at 37°C while shaking at 200 rpm for another 2 h. Co-transformation mixtures were then grown on nonselective (NSM, M9 + His-deficient) and selective (SSM, M9 + His-deficient + 5 mM 3-amino-1,2,4-triazole) screening media for 24-36 h at 37°C. Notably, incubation was conducted for an additional 24-36 h at 30°C in the dark, for the weak interactors. Colonies that grew on both the NSM and SSM plates were identified as positive interaction pairs. However, colonies that were observed only on NSM but not on SSM indicated that there was no interaction between the proteins.

### Pull-down assay

The *in vitro* pull-down assay to assess the protein-protein interaction between YcgR and SpeA was conducted as previous described by using the Pierce GST Protein Interaction Pull-Down Kit (Thermo Fisher)(68). In brief, GST-YcgR was used as the bait, and SpeA-His was used as the prey. For the binding assays, 100 to 150 μg of GST-YcgR was immobilized into 50 μL Glutathione Agarose resin, then the same molar ratio of prey protein SpeA-His was added to the same column for capture. After washed with wash solution, protein complexes were eluted with 10 mM reduced glutathione and analyzed by 10% SDS-PAGE gel.

### RNA isolation, cDNA synthesis, and qRT-PCR analysis

When the bacterial culture was grown to an OD_600_ of target value (0.5, 1.0 and 1.5) in LB medium, bacterial cells were harvested, and total RNA were isolated using Promega Eastep Super Total RNA Extraction Kit according its instruction manual.RNA quality and integrity was assessed by using the NanoDrop 2000 system (Thermo Fisher Scientifific, USA) and agarose gel electrophoresis.

For cDNA synthesis, 1 µg total RNA of each sample was used as the template to synthesize cDNA using the HiScript Ⅲ RT SuperMix for qPCR (+gDNA wiper) (Vazyme, China). ChamQ Universal SYBR qPCR Master Mix (Vazyme, China) was used for qRT-PCR to quantify the transcript levels of the genes of c-di-GMP system and putrescine pathway-related genes, with *gapA* used as the internal reference gene according to MIQE guidelines(69, 70). Primers for SYBR green qRT-PCR were designed using the Beacon designer (Premier Biosoft) and are listed in Table S2. Assays were performed in quadruplicate in 20-µl reaction mixtures using the Applied Biosystems QuantStudio 6 Flex real-time PCR system (Thermo Fisher Scientific, USA). PCR was performed according to the manufacturer’s instructions. The gene expression level was analyzed and calculated using ΔΔ*C*_T_ methods as previously described(71). Data represent the means from three independent biological repeats, with error bars indicating the standard deviations.

### Bacterial growth analysis

Bacterial cultures grown overnight in LB medium were inoculated in the same medium at a dilution of 1:100 and adjusted to the same cell density. 200 µl of the diluted culture was added to each well and grown at 28°C with low intensity shaking, and growth curve was analyzed by the Bioscreen-C automated growth curve analysis system (OY Growth Curves AB, Ltd., Finland).

### Data availability

The genome sequence of *D. oryzea* EC1 is accessible in the NCBI database under accession no. NZ_CP006929.1. The nucleotide sequences of *wspR* and *rocR* in *P. aeruginosa* PAO1 are accessible in the NCBI database under gene IDs 878337 and 878871.

## SUPPLEMENTAL MATERIAL

### Supplemental material is available online only

**FIGURE S1 Swimming motility of deletion mutants of putrescine pathway-associated proteins capable of interacting with YcgR.**

Four putrescine pathway-related proteins that can directly interact with YcgR have been identified in this experiment as follows: SpeA, biosynthetic arginine decarboxylase; ArtP, arginine ABC transporter ATP-binding protein; ArgG, argininosuccinate synthase; MetK, S-adenosylmethionine synthase. Dotted lines indicate the diameter of swimming zone of wild-type EC1. Experiments were repeated three times in triplicates. ****, *P* < 0.0001; ***, *P* < 0.001; ns, *P* > 0.05(by one-way ANOVA with multiple comparisons).

**FIGURE S2 Swimming motility of Δ*ycgR* and the complemented strain Δ*ycgR* (*ycgR*).**

Dotted lines indicate the diameter of swimming zone of wild-type EC1. Experiments were repeated three times in triplicates. ****, *P* < 0.0001; ns, *P* > 0.05(by one-way ANOVA with multiple comparisons).

**FIGURE S3 Deletion of *ycgR* gene in the background of Δ*speA* did not alter the substantially impaired swimming motility and putrescine production of Δ*speA*.**

(A) Swimming motility. Dotted lines indicate the diameter of swimming zone of Δ*speA*. ****, *P* < 0.0001; **, *P* < 0.01(by one-way ANOVA with multiple comparisons). (B) Putrescine production. Dotted lines indicate the putrescine levels of Δ*speA*. *, *P* < 0.05; ns, *P* > 0.05(by Student’s unpaired *t* test). Experiments were repeated three times in triplicates.

**FIGURE S4 *In vitro* overexpressed of *ycgR* in EC1 did not increase production of putrescine but significantly decreased its swimming motility.**

(A) Putrescine production. Dotted lines indicate the putrescine levels of wild-type EC1. ns, *P* > 0.05(by Student’s unpaired *t* test). Experiments were repeated three times in triplicates. (B) Swimming motility. ****, *P* < 0.0001(by Student’s unpaired *t* test). Experiments were repeated three times in triplicates.

**FIGURE S5 Effect of adding different molar ratios of YcgR on the activity of SpeA *in vitro*.**

The product agmatine was extracted after 1h and 2h respectively of the reaction and its absorbance value was measured after 8h. Dotted lines indicate the agmatine production of the system when only SpeA was added. *, *P* < 0.05, ns, *P* > 0.05(by Student’s unpaired *t* test).

**FIGURE S6 YcgR cannot interact with bacterial flagellar-associated proteins.** (A) Bacterial two-hybrid system analysis(YcgR1 and FilG1 belongs to EC1; YcgR2 and FilG2 belongs to *E. coli*). (B) Pull-down analysis(YcgR and FilG belongs to EC1). (C) Bacterial two-hybrid system analysis of YcgREC1 and other flagellar structure proteins.

**FIGURE S7 Time-course analysis of *ycgR* and *speA* gene expression in wild-type strain EC1 by RT-qPCR.** Total RNAs were collected when bacterial cells were grown to the corresponding OD_600_.

TABLE S1 Strains and plasmids used in this study.

TABLE S2 Primers used in this study.

## FOUNDINGS

This study was supported by grants from the Key Realm R&D Program of Guangdong Province (2020B0202090001), National Natural Science Foundation of China (31900076, 32102155 and 31901843) and Natural Science Foundation of Guangdong Province (2020A1515110022, 2022A1515010564).

W.G., Y.C., L.L performed experiments. W.G., Y.C., L.L and L.Z designed experiments, analyzed the data, and wrote the paper.

We declare that there are no known conflicts of interest associated with this paper.

## ACKNOWLEDGMENTS

We thank the Shanghai Applied Protein Technology Co. Ltd. for technology support in IP-MS analysis.

