## Supplementary material for "Cyclic di-GMP interact with putrescine via a PilZ domain receptor YcgR": Table S1-S2

**Table S1. Strains and plasmids used in this study.**

| Strain or plasmid | Relevant characteristics[a] | References of source |
|---|---|---|
| **Strains** | | |
| *Dickeya oryzae* | | |
| EC1 | Wild-type strain, Pm$^r$ | Lab collection |
| 7ΔPDE | Deletion of all c-di-GMP degradation genes of EC1, Pm$^r$ | Lab collection |
| 7ΔPDEΔ*ycgR* | *W909_08750* gene deletion of 7ΔPDE, Pm$^r$ | Lab collection |
| Δ*ycgR* | *W909_08750* gene deletion mutant, Pm$^r$ | Lab collection |
| 15ΔDGC | Deletion of all c-di-GMP synthase genes of EC1, Pm$^r$ | Lab collection |
| Δ*speA* | *W909_17465* gene deletion mutant, Pm$^r$ | This study |
| Δ*argG* | *W909_00315* gene deletion mutant, Pm$^r$ | This study |
| Δ*artP* | *W909_08445* gene deletion mutant, Pm$^r$ | This study |
| Δ*metK* | *W909_17475* gene deletion mutant, Pm$^r$ | This study |
| Δ*speA*Δ*ycgR* | *W909_08750* gene deletion of Δ*speA*, Pm$^r$ | This study |
| Δ*speA*Δ*potF*Δ*plaP* | *W909_08395* and *W909_08760* gene deletion of Δ*speA*, Pm$^r$ | This study |
| 15ΔDGCΔ*speA* | *W909_17465* gene deletion of 15ΔDGC,Pm$^r$ | This study |
| 7ΔPDEΔ*speA* | *W909_17465* gene deletion of 7ΔPDE,Pm$^r$ | This study |
| Δ*ycgR*(*ycgR*) | Δ*ycgR* carrying pBBR1-ycgR vector, Ap$^r$ | This study |
| 7ΔPDE(*rocR*) | 7ΔPDE carrying pBBR1-rocR vector, Ap$^r$ | This study |
| 15ΔDGC(*wspR*) | 15ΔDGC carrying pBBR1-wspR vector, Ap | Lab collection |
| Δ*speA*Δ*potF*Δ*plaP*(*rocR*) | Δ*speA*Δ*potF*Δ*plaP* carrying pBBR1-rocR vector, Ap$^r$ | This study |
| EC1(*ycgR*) | Wild-type strain carrying pBBR1-ycgR vector, Pm$^r$ | This study |
| *Escherichia coli* | | |
| K12 CC118 | *gyrA*， *recA*， *λ pir* | Lab collection |
| DH5α | *deoR*, *recA*, *endA*, *hsdR*, *supE*, *thi*, *gyrA*, *relA* | Lab collection |
| BL21 | $B\ F^{-}\ ompT\ gal\ dcm\ lon\ hsdS_B(r_B^{-}m_B^{-})\ [malB^{+}]_{K\text{-}12}(\lambda^S)$ | Lab collection |
| **Plasmids** | | |
| pKNG101 | Suicide vector; Str$^r$, SacB, mobRK2, oriR6K (pir-minus) | Lab collection |
| pKNG-*ycgR* | *W909_08750* knock-out fragment ligated on pKNG101 | This study |
| pKNG-*speA* | *W909_17465* knock-out fragment ligated on pKNG101 | This study |
| pKNG-*potF* | *W909_08395* knock-out fragment ligated on pKNG101 | This study |
| PKNG-*plaP* | *W909_08760* knock-out fragment ligated on pKNG101 | This study |
| pKNG-*argG* | *W909_00315* knock-out fragment ligated on pKNG101 | This study |
| pKNG-*artP* | *W909_08445* knock-out fragment ligated on pKNG101 | This study |
| pKNG-*metK* | *W909_17475* knock-out fragment ligated on pKNG101 | This study |
| pBBR1MCS-4 | Expression vector contains a *lacZ* promoter, Ap$^r$ | Lab collection |
| pBBR1-*ycgR* | pBBR1-MCS4 carries the coding region of *W909_08750* at down-stream of lac promoter, Ap$^r$ | This study |
| pBBR1-*rocR* | pBBR1-MCS4 carries the coding region of PA3947 at down-stream of lac promoter, Ap$^r$ | Lab collection |

| | | |
|---|---|---|
| pGEX-6P-1 | Protein expression vector containing GST tag, $Ap^r$ | Lab collection |
| pGEX-YcgR | pGEX-6P-1 carries the coding region of *W909_08750* at down-stream of GST tag, $Ap^r$ | This study |
| pET-32a | Protein expression vector containing HIS tag, $Ap^r$ | Lab collection |
| pET-SpeA | pET-32a carries the coding region of *W909_16940* at down-stream of HIS tag, $Ap^r$ | This study |
| pBT | Two-hybrid system bait plasmid containing the *cat* gene, p15A origin of replication and λ cI ORF | Lab collection |
| pBT-YcgR1 | pBT containing *ycgR* of EC1 | This study |
| pBT-YcgR2 | pBT containing *ycgR* of *e.coli* | This study |
| pTRG | Two-hybrid system target plasmid containing the *tet* gene, ColE1 origin of replication, and RNA polymerase α subunit ORF | Lab collection |
| pTRG-FliG1 | pTRG containing *fliG* of EC1 | This study |
| pTRG-FliG2 | pBT containing *fliG* of *e.coli* | This study |
| pTRG-SpeA | pTRG containing *speA* of EC1 | This study |
| pTRG-ArgG | pTRG containing *argG* of EC1 | This study |
| pTRG-ArtP | pTRG containing *artP* of EC1 | This study |
| pTRG-MetK | pTRG containing *metK* of EC1 | This study |

Abbreviations: $Ap^r$, ampicillin resistance; $Pm^r$, polymyxinB resistance; $Kan^r$, kanamycin resistance; $Str^r$, streptomycin resistance;$Cm^r$,chloramphenicol resistance.

**Table S2. Primers used in this study.**

| Primer name | Primer sequence (5'-3') |
|---|---|
| **For deletion** | |
| SpeA-1 | cccctgcaggtcgacggatccCTTTTCTTCTGACGCCACTGC |
| SpeA-2 | ttctaacgagagaGGCGATCCCCTTCTTCCA |
| SpeA-3 | gatcgccTCTCTCGTTAGAAAAGGTTGAGTGC |
| SpeA-4 | cggactatagactatactagtCGGGACAACAACGCCGCC |
| PotF-1 | cccctgcaggtcgacggatccTACGTTATGGATCAATAGCTGGTCA |
| PotF-2 | CCGTTCCTTCCTCCATACCAG |
| PotF-3 | tggtatggaggaaggaacggCCGCTGTCCTTGACTCCACTT |
| PotF-4 | cggactatagactatactagtCTGCACAAATTTACCGCGATT |
| PlaP-1 | cccctgcaggtcgacggatccTGGCTAGCTGGGGCAAAAT |
| PlaP-2 | cctgataTACGCAAACCTCCTTTACCGA |
| PlaP-3 | ggaggtttgcgtaTATCAGGTGATGAGAACGGGCT |
| PlaP-4 | cggactatagactatactagtAGATATTCAGCCACTACTGGCGC |
| ArgG-1 | cccctgcaggtcgacggatccGCGGTGGTGGGGGGATAT |
| ArgG-2 | caggacgcgcCATAACTATTAGTCCCTGCTTGATTTC |
| ArgG-3 | aatagttatgGCGCGTCCTGAAGGACATG |
| ArgG-4 | cggactatagactatactagtCGTGATATTCATAAGTAACACTATATATTACGA |
| ArtP-1 | cccctgcaggtcgacggatccGTTGTTCGAGGCAGCCTATTATT |
| ArtP-2 | gcatgcgtaCATCAAACCGTCCTTCTCTTCAA |

| | |
|---|---|
| ArtP-3 | acggtttgatgTACGCATGCCTCACCGTCC |
| ArtP-4 | cggactatagactatactagtCGGTGTTAGTGGCGAAGAACG |
| MetK-1 | cccctgcaggtcgacggatccAATCCTATCACCTCCACGCTCG |
| MetK-2 | catgtcagaaaagcggaGAGTTTCTTTACCTTATAAGCTAAAGCC |
| MetK-3 | ctcTCCGCTTTTCTGACATGCCA |
| MetK-4 | cggactatagactatactagtATTCCAACGTCACCTGCAGG |
| YcgR-1 | cccctgcaggtcgacggatccTTTCACACTATCCTGCCGCTT |
| YcgR-2 | tatctgtttgacttcgcCCCCACCGTCGATTGAATG |
| YcgR-3 | gggGCGAAGTCAAACAGATAGAGCGC |
| YcgR-4 | cggactatagactatactagtAGTCCCATCATTACCACTGGCA |
| PKNG-F | GCCATCAAACCACGTCAAAT |
| PKNG-R | AACCAAGCCTATGCCTACAG |
| **For *in trans* expression** | |
| CYcgR-1 | gataagcttgatatcgaattcATGGATGTAGTGGATGACAATATGAAA |
| CYcgR-2 | cgctctagaactagtggatccATAGTACAGCACGTCAGGTATCGC |
| MCS-F | TCTTCGCTATTACGCCAGCT |
| MCS-R | GGCTCGTATGTTGTGTGGAA |
| **For protein expression** | |
| pGEX-(ycgR)-1 | ttccaggggcccctgggatccATGGATGTAGTGGATGACAATATGAAA |
| pGEX-(ycgR)-2 | ctcgagtcgacccgggaattcAACCGACGCGATAACGGTG |
| pGEX-F | GGGCTGGCAAGCCACGTTTGGTG |
| pGEX-R | CCGGGAGCTGCATGTGTCAGAGG |

| | |
|---|---|
| pET-(speA)-1 | gccatggctgatatcggatccATGTCTGACGATATGATTCAACCG |
| pET-(speA)-2 | ctcgagtgcggccgcaagcttCTCGTCTTCCAGATAAGTGTAACCG |
| pET-F | CATTCTTCTGGTCTGGTG |
| pET-R | CTCAAGACCCGTTTAGAG |
| **For bacterial two-hybrid assay** | |
| pB-ycgR-1(e.coli) | tggcgcggccgcatcgaattcCGTGAGTCATTACCATGAGCAGTT |
| pB-ycgR-2(e.coli) | aattaattaactcgaggatccTCAGTCGCGCACTTTGTCC |
| pB-ycgR-1(EC1) | tggcgcggccgcatcgaattcGATGGATGTAGTGGATGACAATATGA |
| pB-ycgR-2(EC1) | aattaattaactcgaggatccTCAACGCAGGCGTTTGCG |
| pT-fliG-1(e.coli) | gatccgcggccgcaagaattcTCATGAGTAACCTGACAGGCACC |
| pT-fliG-2(e.coli) | ttaattaattaattactcgagTCAGACATAGGTATCCTCGCCG |
| pT-speA-1(EC1) | gatccgcggccgcaagaattcCCATGTCTGACGATATGATTCAACC |
| pT-speA-2(EC1) | ttaattaattaattactcgagCTCGTCTTCCAGATAAGTGTAACCG |
| pT-argG-1(EC1) | gatccgcggccgcaagaattcTTATGACGACGATTTTGAAACATCT |
| pT-argG-2(EC1) | ttaattaattaattactcgagTTTCGCGGACTTATCCGGC |
| pT-artP-1(EC1) | gatccgcggccgcaagaattcTGATGATTTTCCTGAAAAATGTTTCT |
| pT-artP-2(EC1) | ttaattaattaattactcgagATGCAGGATTTTAGCCAAAAAGTC |
| pT-metK-1(EC1) | gatccgcggccgcaagaattcTCATGGCTAAACACCTTTTTACATC |
| pT-metK-2(EC1) | ttaattaattaattactcgagTTTCAGGCCTGCGGCATC |
| pBT-F | TCCGTTGTGGGGAAAGTTATC |
| pBT-R | GGGTAGCCAGCAGCATCC |
| pTRG-F | TGGCTGAACAACTGGAAGCT |

| | |
|---|---|
| pTRG-R | ATTCGTCGCCCGCCATAA |
| **For qPCR** | |
| DdgcA-Q1 | TATGTCGCTGATAAGATTC |
| DdgcA-Q2 | CGTTAATGGCTAATGTCA |
| DpdeA-Q1 | TTCATTATGCTGCTACAA |
| DpdeA-Q2 | TACAGTAAGTGGCTATCA |
| DpdeB-Q1 | ACTGCTATCGTTTCTTTA |
| DpdeB-Q2 | TTGTTATTGACCATATTGAC |
| speA-Q1 | CGATATGCTGGAATATGT |
| speA-Q2 | TAAGTGTAACCGTATAACC |
| speC-Q1 | GACGGCACTATCTATAAC |
| speC-Q2 | GATGAACTGCTCATATCC |
| metK-Q1 | CGAAGGACATCCTGATAA |
| metK-Q2 | CACCAACTAACACCATAC |
| artP-Q1 | AATGTGAACAAGTGGTAT |
| artP-Q2 | GGATTGTGGTTGATTTAC |
| argG-Q1 | GACAACCTGACATACAAA |
| argG-Q2 | ATATCCAGATTACGCATC |
| YcgR(qPCR)-1 | TAGTGGATGACAATATGA |
| YcgR(qPCR)-2 | AGGATCTTACTGATGAAT |
