## Supplementary figures and images for "Cyclic di-GMP interact with putrescine via a PilZ domain receptor YcgR"

### Figure S1-S7

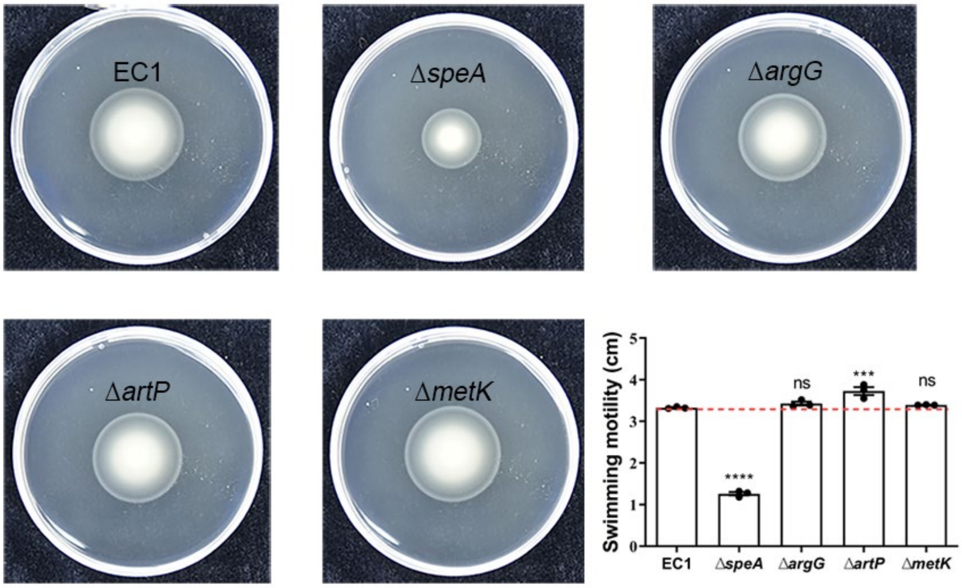

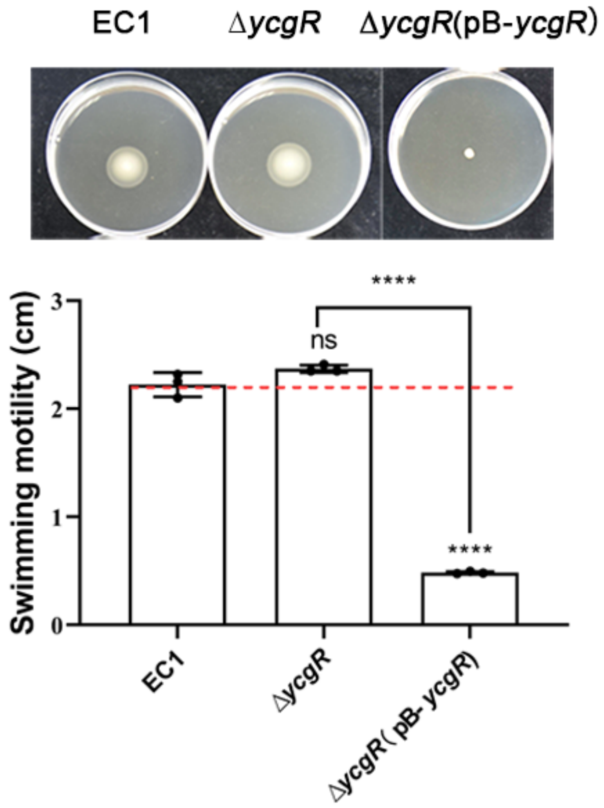

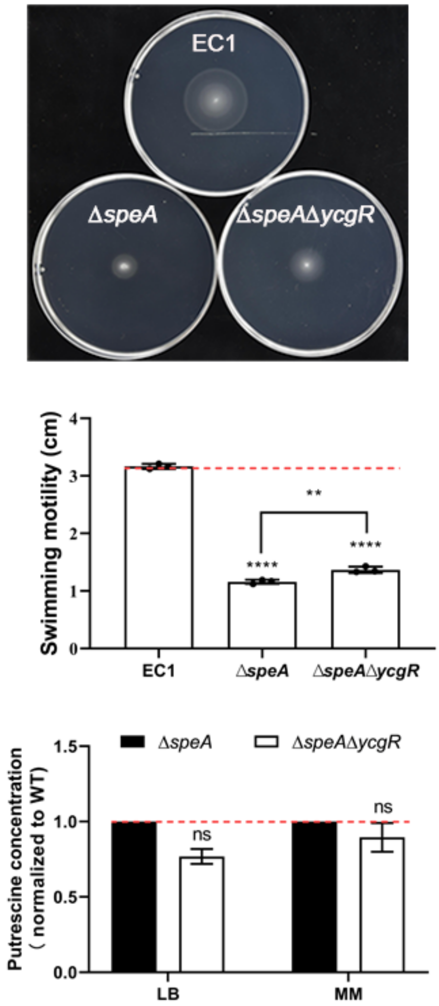

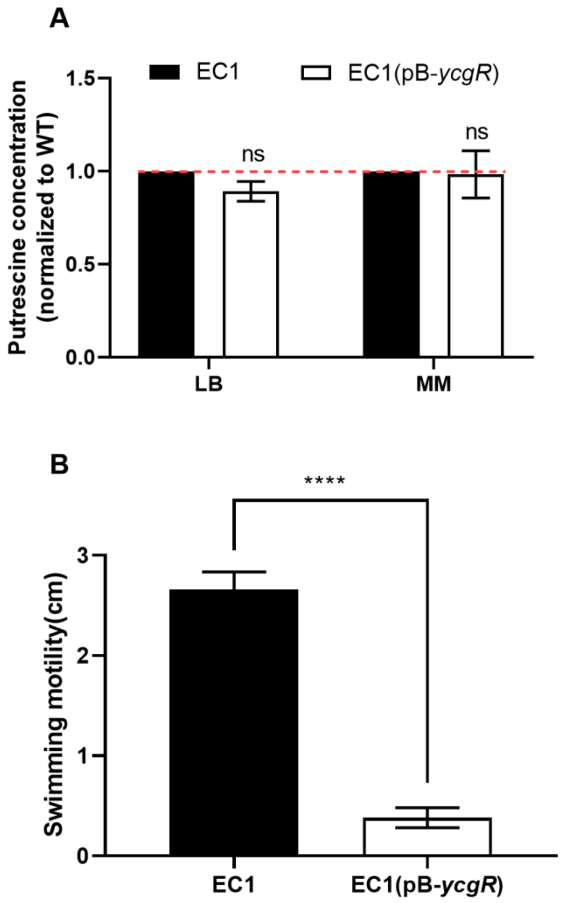

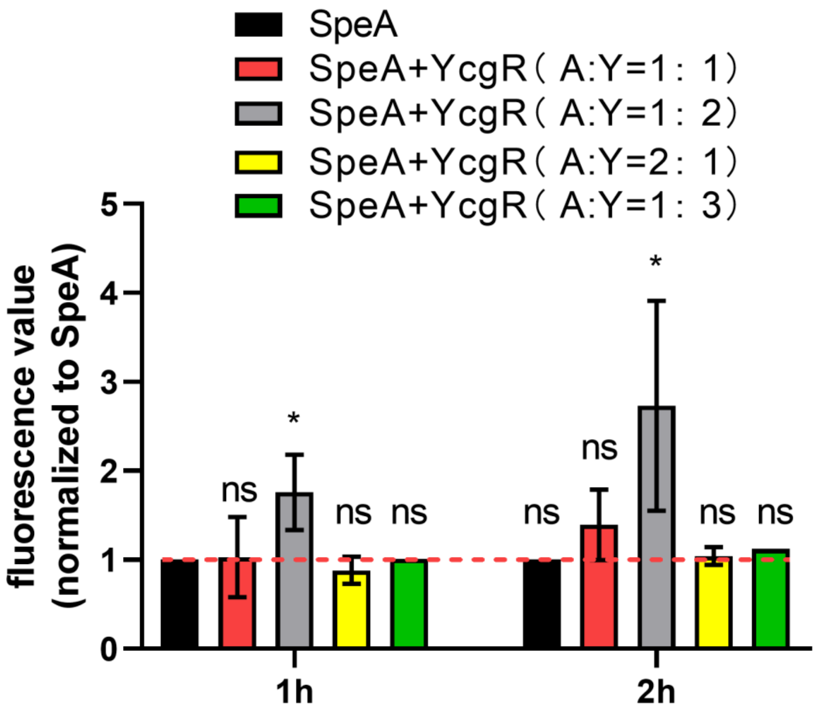

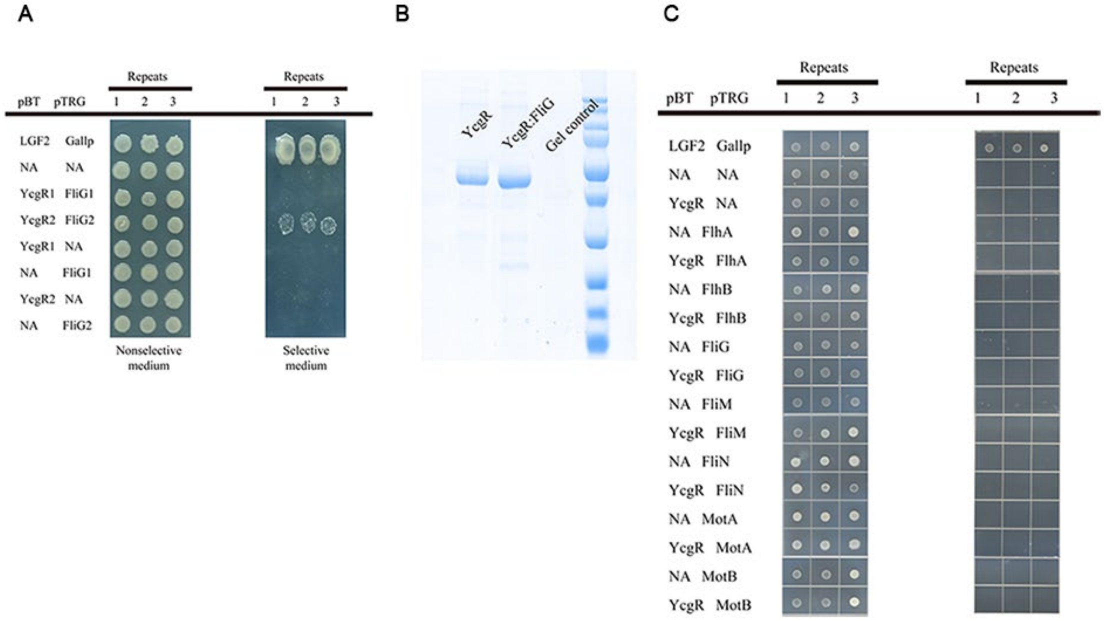

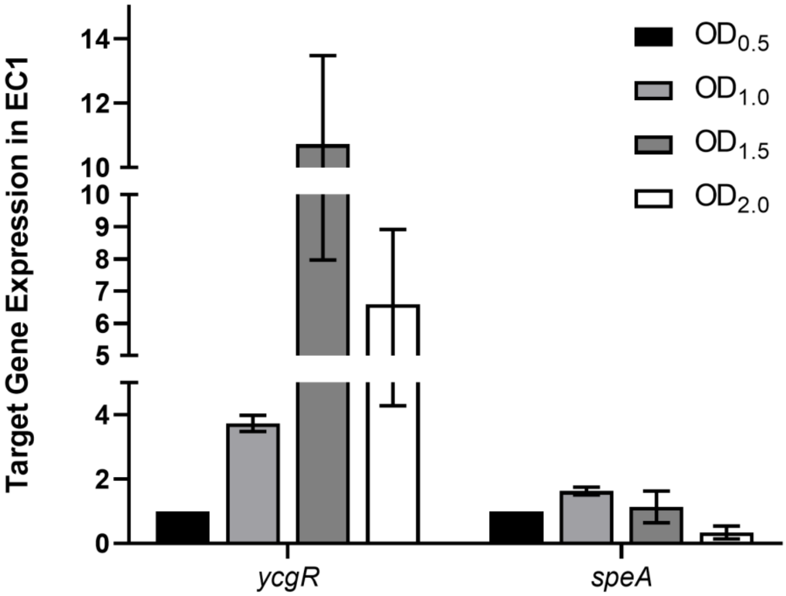
